# Functional prediction and comparative population analysis of variants in genes for proteases and innate immunity related to SARS-CoV-2 infection

**DOI:** 10.1101/2020.05.13.093690

**Authors:** Kristel Klaassen, Biljana Stankovic, Branka Zukic, Nikola Kotur, Vladimir Gasic, Sonja Pavlovic, Maja Stojiljkovic

**Author notes:** Corresponding author (MS). These authors contributed equally to this work (Kristel Klaassen, Biljana Stankovic).

## Abstract

New coronavirus SARS-CoV-2 is capable to infect humans and cause a novel disease COVID-19. Aiming to understand a host genetic component of COVID-19, we focused on variants in genes encoding proteases and genes involved in innate immunity that could be important for susceptibility and resistance to SARS-CoV-2 infection.

Analysis of sequence data of coding regions of *FURIN, PLG, PRSS1, TMPRSS11a, MBL2* and *OAS1* genes in 143 unrelated individuals from Serbian population identified 22 variants with potential functional effect. *In silico* analyses (PolyPhen-2, SIFT, MutPred2 and Swiss-Pdb Viewer) predicted that 10 variants could impact the structure and/or function of proteins. These protein-altering variants (p.Gly146Ser in *FURIN*; p.Arg261His and p.Ala494Val in *PLG*; p.Asn54Lys in *PRSS1*; p.Arg52Cys, p.Gly54Asp and p.Gly57Glu in *MBL2*; p.Arg47Gln, p.Ile99Val and p.Arg130His in *OAS1*) may have predictive value for inter-individual differences in the response to the SARS-CoV-2 infection.

Next, we performed comparative population analysis for the same variants using extracted data from the 1000 genomes project. Population genetic variability was assessed using delta MAF and Fst statistics. Our study pointed to 7 variants in *PLG, TMPRSS11a, MBL2* and *OAS1* genes with noticeable divergence in allelic frequencies between populations worldwide. Three of them, all in *MBL2* gene, were predicted to be damaging, making them the most promising population-specific markers related to SARS-CoV-2 infection.

Comparing allelic frequencies between Serbian and other populations, we found that the highest level of genetic divergence related to selected loci was observed with African, followed by East Asian, Central and South American and South Asian populations. When compared with European populations, the highest divergence was observed with Italian population.

In conclusion, we identified 4 variants in genes encoding proteases (*FURIN, PLG* and *PRSS1*) and 6 in genes involved in the innate immunity (*MBL2* and *OAS1*) that might be relevant for the host response to SARS-CoV-2 infection.

## Introduction

A new coronavirus SARS-CoV-2 capable to infect humans, emerged in mid-December 2019 in Wuhan, China, causing the novel disease COVID-19 [1]. Clinical manifestations of this viral infection vary from asymptomatic to severe acute respiratory syndrome and death. The number of infected people and affected countries have quickly risen and already in March 2020, World Health Organization declared a pandemic [2]. Therefore, it became highly important to understand if specific human genes and genetic variants could be associated with susceptibility or resistance to SARS-CoV-2 infection and how frequencies of these variants vary between different populations. Human genes usually associated with susceptibility and resistance to viral infection are those associated with the point of viral entry into the human host cells, such as genes encoding receptors, co-receptors and enzymes that modify receptors [3]. Furthermore, various genes involved in the immune response, such as virus sensing, signaling in response to virus, antiviral factors etc. have also been found to be important for the severity and the outcome of viral infections. A recent study which analyzed COVID-19 symptoms in monozygotic and dizygotic twins, reported that 50 (95% confidence intervals 29-70)% of the variance of ‘predicted COVID-19’ phenotype is due to genetic factors [4].

Deciphering the RNA sequence of the SARS-CoV-2 genome showed that this new virus belongs to lineage B betacoronaviruses, together with the SARS-CoV virus which emerged in 2002 [1]. Genetic susceptibility and resistance to SARS-CoV had been extensively studied by genotyping SARS patients with extremely severe and extremely mild clinical manifestations. As a result, variants in *OAS1, MX1, MBL2, CCL2, CCL5, ASHG, IFNgamma, CD14* and *CD209* genes were associated with genetic susceptibility to SARS-CoV [5–11]. Majority of variants that emerged in such studies were located in non-coding parts of the human genome and their effect is observable in the fine tuning of the gene expression. Also, variants residing in coding regions, such as rs1800450 in *MBL2* gene [6] were particularly interesting as they alter protein structure and function.

SARS-CoV and SARS-CoV-2 share ∼76% amino acid sequence identity of the spike (S) protein sequence, a crucial part of the viral envelope which enables specific binding to the receptors at human cells, therefore contributing to viral potential to infect humans [12, 13]. It was shown that SARS-CoV-2 binds to the human angiotensin-converting enzyme 2 (ACE2) receptors, as SARS-CoV does, however with the higher affinity [14]. Human transmembrane protease serine 2 (TMPRSS2), an enzyme important for the entry of SARS-CoV [15], was recently found to activate the S protein of the SARS-CoV-2 [16].

Moreover, it was found that the S protein of the SARS-CoV-2 contains a furin-like cleavage site which is absent in coronaviruses of the same clade [17]. Knowing that furin cleavage sites are responsible for the high virulence of human influenza viruses [18], it was suggested that furin-like site in the S protein of SARS-CoV-2 represents its advantage in attaching to the human cells expressing ACE2 receptor.

The SARS-CoV-2 relies on the host cell proteases to cut its S protein in two parts thus forming N-terminal part which recognizes ACE2 receptor and C-terminal part involved in the viral entry which must be further cleaved by furin and/or other furin-like enzymes [17]. It has been shown that plasmin is also capable to cleave furin sites [19] and that individuals with elevated plasmin demonstrated higher susceptibility to COVID-19 [20]. In addition to plasmin, S protein of coronaviruses may be cleaved by other airway proteases such as trypsin-1 and TMPRSS11a [20].

At the beginning of May 2020, in the full swing of COVID-19 pandemic, the data on genetic susceptibility or resistance to SARS-CoV-2 which would be based on the genotyping of individuals infected by SARS-CoV-2 are still lacking. However, a detailed study using computational prediction methods for protein structure analyses revealed that 17 variants in the coding regions of the *ACE2* gene are located at positions important for the binding of ACE2 with the SARS-CoV-2 S protein [21]. Based on these predictions, individuals carrying these variants would probably be resistant to SARS-CoV-2 infection. Yet, all these variants were rare (less than 0.00388 allele frequency) and interestingly enough their association with any human disease or disorder has never been reported [21]. Having in mind the rarity of variants that could directly influence the binding of SARS-CoV-2 to ACE2, it is not surprising that comparative genetic analysis of the frequency of *ACE2* variants in different populations did not predict the existence of individuals resistant to SARS-CoV-2 infection [22].

ACE2 is not the only possible player that could influence the interaction between humans and SARS-CoV-2 virus. The furin could be the second important factor to contribute to high pathogenicity of the novel virus. Furthermore, we can learn from clinical experiences of the SARS-CoV-2 infection which showed that plasmin and other airway proteases could also be important for patients experiencing more severe form of COVID-19 [20]. In addition, the knowledge based on SARS epidemic shows that *MBL2* and *OAS1* genes, involved in the innate immune response, could also modulate susceptibility to infection with betacoronaviruses.

Taking all these players into account, we performed genetic analysis of variants in coding regions of *FURIN*, plasminogen (*PLG*), trypsin-1 (*PRSS1*), *TMPRSS11a*, *MBL2* and *OAS1* genes in Serbian populations aiming to identify possible genetic markers that are capable to impact protein structure/function and thus contribute to the susceptibility or resistance to SARS-CoV-2 infection.

We postulate that the variants in genes encoding proteases could be advantageous, while variants in genes encoding proteins involved in the innate immunity add some disadvantage to individuals in combating COVID-19. We also performed comparative genetic analysis in different populations in order to assess the divergence between populations for those variants.

## Subjects and methods

### Genetic and bioinformatic analysis

In this study, we analyzed the genomic sequence data of unrelated individuals from Serbian population, extracted from the in-house database of Laboratory for Molecular Biomedicine, Institute of Molecular Genetics and Genetic Engineering, University of Belgrade. Written informed consent was obtained from all participants. The study was conducted in accordance with the Helsinki Declaration and approved by the Ethic Committee of Institute of Molecular Genetics and Genetic Engineering, University of Belgrade.

Total of 143 unrelated Serbian individuals (84 males and 59 females) were previously analyzed by NGS approach using the Illumina Clinical Exome Sequencing TruSight One Gene Panel (Illumina, San Diego, CA, USA), as previoustly described [23]. VCF files were further annotated and examined using the Illumina VariantStudio 3.0 Data Analysis Software (Illumina, San Diego, CA, USA). Variants that did not pass variant call quality filters (those with quality score <Q20, read depth <20, percentage of variant frequency for the minor allele <20% and homopolymer length >8) in more than 10% of samples were excluded. Variants that were considered for further analysis were either missense, start loss, stop gain, splice region variants and frameshift, while synonymous variants were omitted. For all selected variants, Hardy-Weinberg equilibrium was tested, using the chi square goodness of fit test.

Next, we compared genotype data from Serbian population with European (Italy, Spain, Finland, Great Britain and USA with European ancestry), as well as other 4 super-populations, Eastern Asians, South Asians, African and Ad Mixed American - Central and South American populations (total of 2504 subjects). Genotype data were extracted from the VCF files of Phase 3 variant calls of the 1000 Genomes Project (1kGP) sample collection (https://www.internationalgenome.org/) via Ensembl Data Slicer Tool. Populations and details regarding the 1kGP data were described previously [24]. Fisher exact test was used to measure significant differences in genotypes distributions between Serbian and 1kGP populations, applying Bonferoni correction for multiple testing.

We examined the level of population genetic variability at each selected loci using: (1) the maximal global differences in minor allele frequencies (delta MAF) calculated by subtracting the maximum and the minimum MAF across analyzed population groups, (2) using Fst statistics [25], which is widely used in population genetics [26].

R software was utilized for genotype data manipulation, statistical calculations as well as graphical presentations. For estimating Fst statistics, R packages adegenet and hierfstat were used.

### *In silico* prediction analysis

Protein sequences were downloaded from Ensemble, accessed on 2020/04/24 at http://www.ensembl.org. To predict the effect of nonsynonymous amino acid substitutions, we used *in silico* prediction algorithms: PolyPhen-2 (http://genetics.bwh.harvard.edu/pph2), SIFT/PROVEAN (http://provean.jcvi.org/index.php) and MutPred2 (http://mutpred.mutdb.org/). The Swiss-Pdb Viewer (Swiss Institute of Bioinformatics, at http://www.expasy.org/spdbv) was used to analyze the effects of variants upon the structure of the proteins, using the crystal structures with PDB codes 4ig8.pdb for the structure of human OAS1 in complex with dsRNA and 2’-deoxy ATP, and 4z2a.pdb for the structure of unglycosylated apo human furin (Protein Data Bank-RCSB, accessed on the 2020/04/24, at http://www.rcsb.org/pdb).

## Results

### Analysis of genetic variants found in Serbian population

Analysis of sequence data of coding regions of *FURIN, PLG, PRSS1, TMPRSS11a, MBL2* and *OAS1* genes in 143 unrelated individuals from Serbian population identified 22 variants with potential functional effects (Table 1). Among identified variants, there were 19 missense variants, 1 start lost, 1 missense/splice region and 1 splice region variant. Majority of the detected variants were rare (8 variants had minor allele frequency of 0.3%, while 4 variants of 0.7%). We detected the highest minor allelic frequency for the *OAS1* p.Gly162Ser, being 41%. All detected variants were in Hardy-Weinberg equilibrium.

**Table 1.**
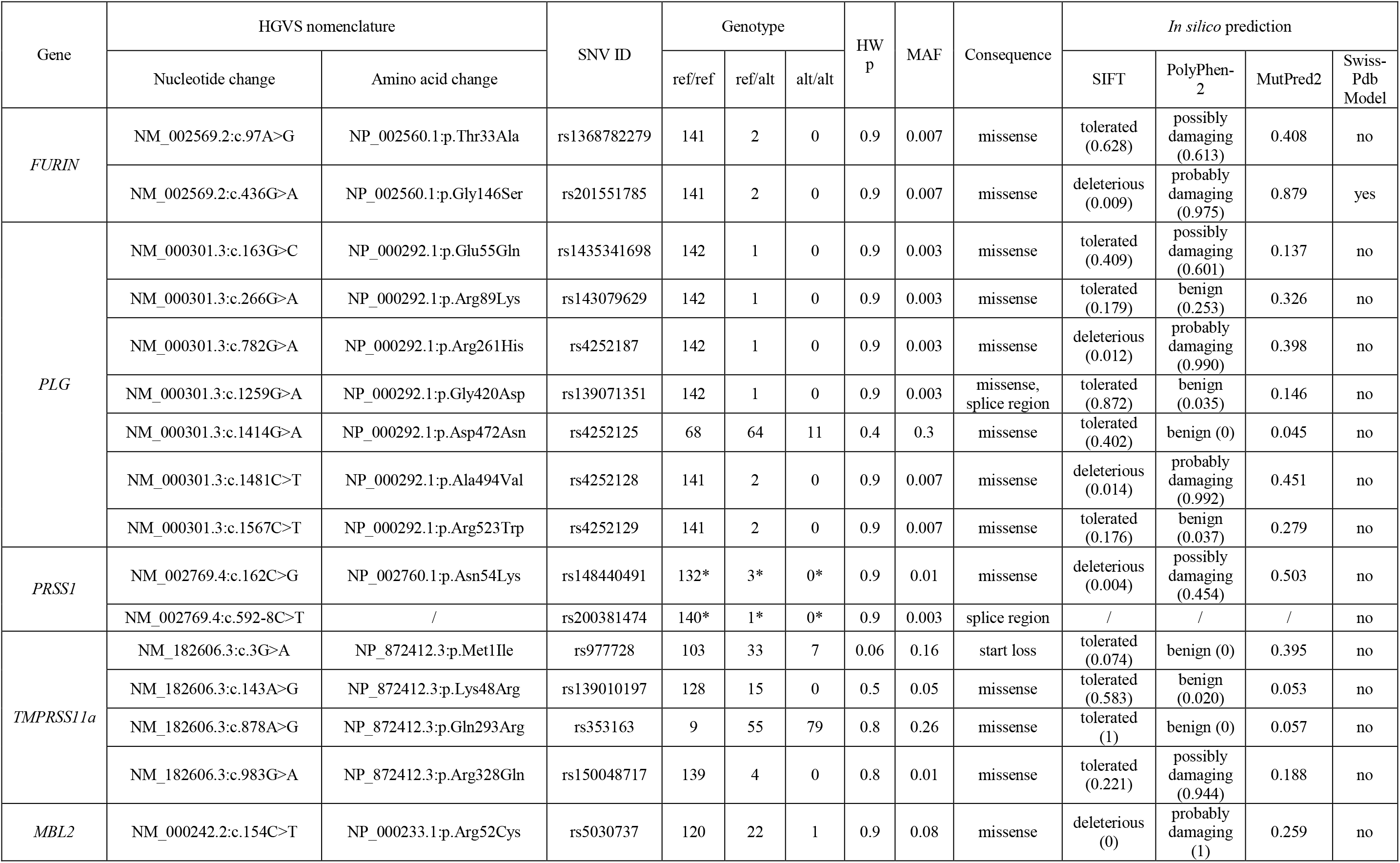
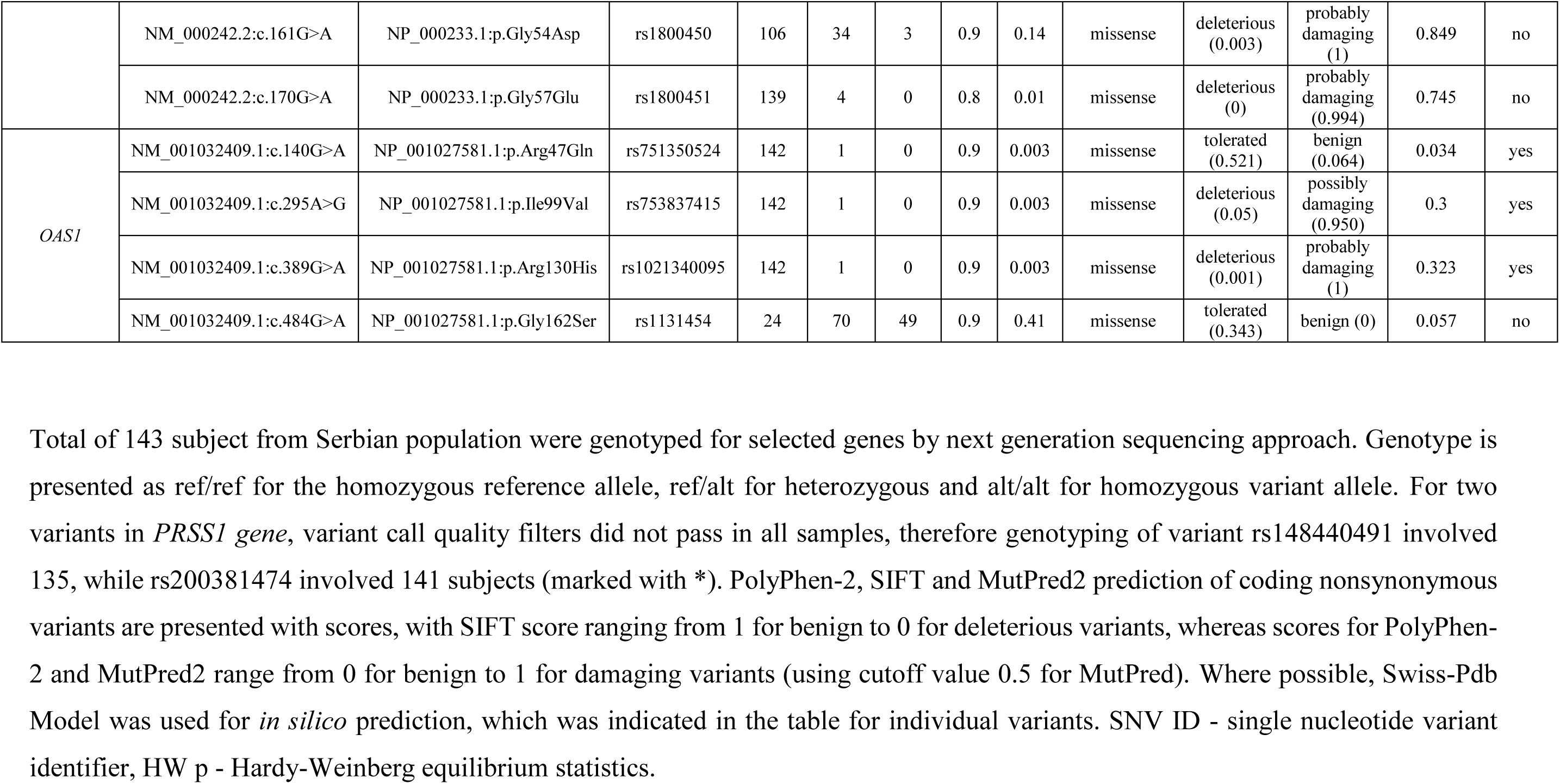
Genetic variants detected in Serbian population.

### *In silico* prediction analysis

Next, we performed *in silico* prediction analysis of the identified variants’ effect, using PolyPhen-2, SIFT and MutPred2 algorithms (Table 1). Nine missense variants were predicted to be deleterious/damaging by both PolyPhen-2 and SIFT (p.Gly146Ser in *FURIN;* p.Arg261His and p.Ala494Val in *PLG;* p.Asn54Lys in *PRSS1;* p.Arg52Cys, p.Gly54Asp and p.Gly57Glu in *MBL2;* p.Ile99Val and p.Arg130His in *OAS1*), while MutPred2 provided additional indication of pathogenicity for 4 of them (p.Gly146Ser in *FURIN*, p.Asn54Lys in *PRSS1*, p.Gly54Asp and p.Gly57Glu in *MBL2*). Although variant p.Arg47Gln in *OAS1* is predicted to be benign/tolerated by PolyPhen-2, SIFT and MutPred2 algorithms, the analysis in Swiss-Pdb Viewer showed that the side chain of Arg47, unlike the glutamine at this position, was predicted to be important for dsRNA binding.

Among those 10 genetic variants predicted to impact the structure and/or function of proteins, 8 were found to be rare (p.Gly146Ser in *FURIN*; p.Arg261His and p.Ala494Val in *PLG*; p.Asn54Lys in *PRSS1*; p.Gly57Glu in *MBL2*; p.Arg47Gln, p.Ile99Val and p.Arg130His in *OAS1*). These variants may have predictive value for inter-individual differences in the response to the SARS-CoV-2 infection.

Other 2 variants with potentially damaging effect, p.Arg52Cys and p.Gly54Glu in *MBL2* gene, having allelic frequency of 8% and 14%, respectively, are the most promising population-specific markers to be considered for association study in COVID-19 patients in Serbia.

In total, 4 variants in genes encoding proteases (*FURIN, PLG* and *PRSS1*) and 6 in genes involved in the innate immunity (*MBL2* and *OAS1*) might be interesting for further studies in relation to the response to SARS-CoV-2 infection.

### Comparative population analysis

Our next goal was to investigate the allele frequency in populations worldwide for the same 22 genetic variants detected in Serbian population. The minor allele frequencies (MAF) were extracted from the 1kG project involving European populations (Italy, Spain, Finland, Great Britain and USA with European ancestry) as well as Eastern Asians, South Asians, African and Ad Mixed American (Central and South American populations) (Table 2). We denoted MAF as the frequency of the minor allele found in European populations. Majority of variants were with low frequency among all populations (MAF≤ 0.05). From all detected variants, 17 were shared between at least two populations, while 5 were found only in Serbian population.

**Table 2.**
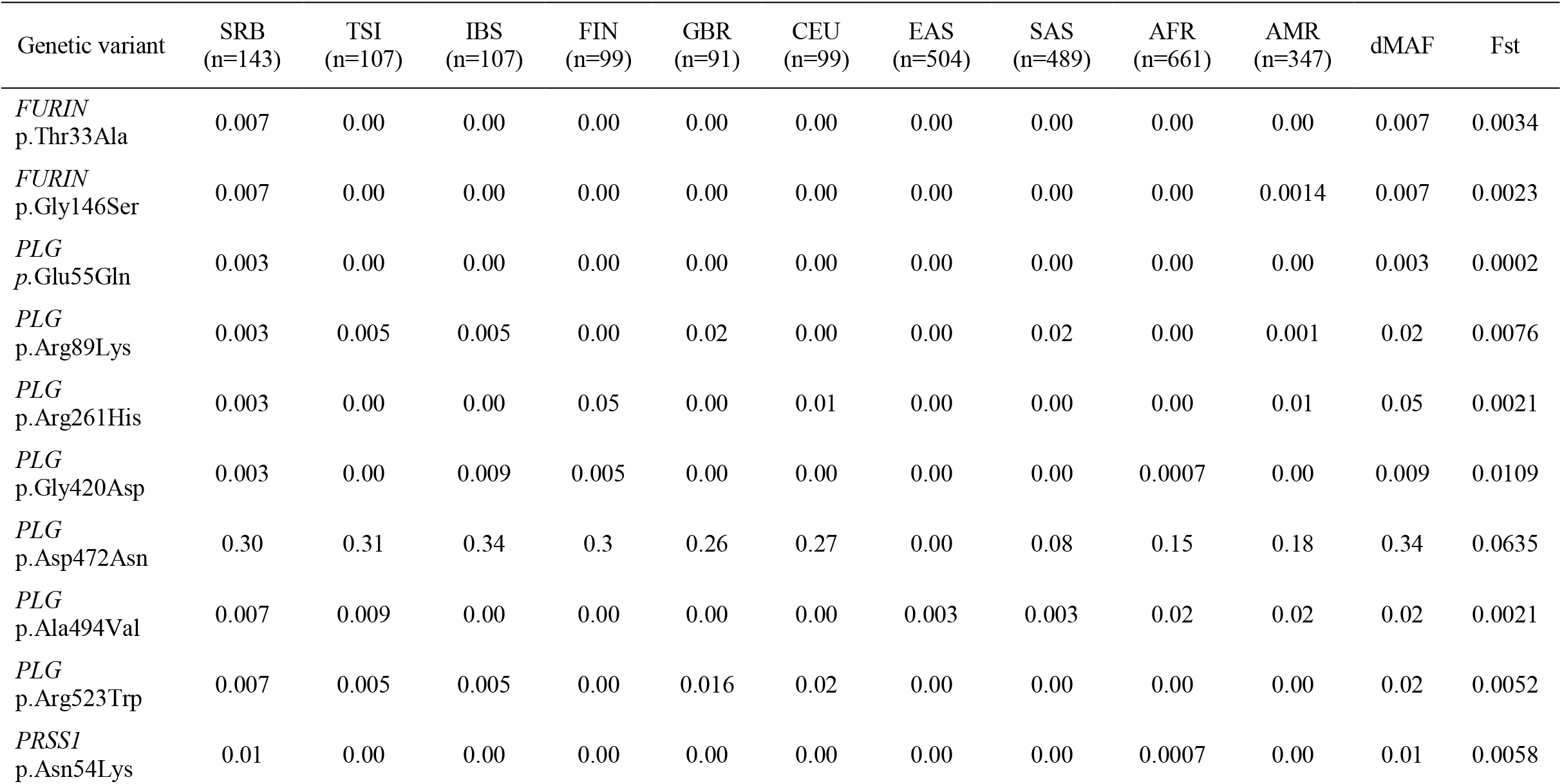
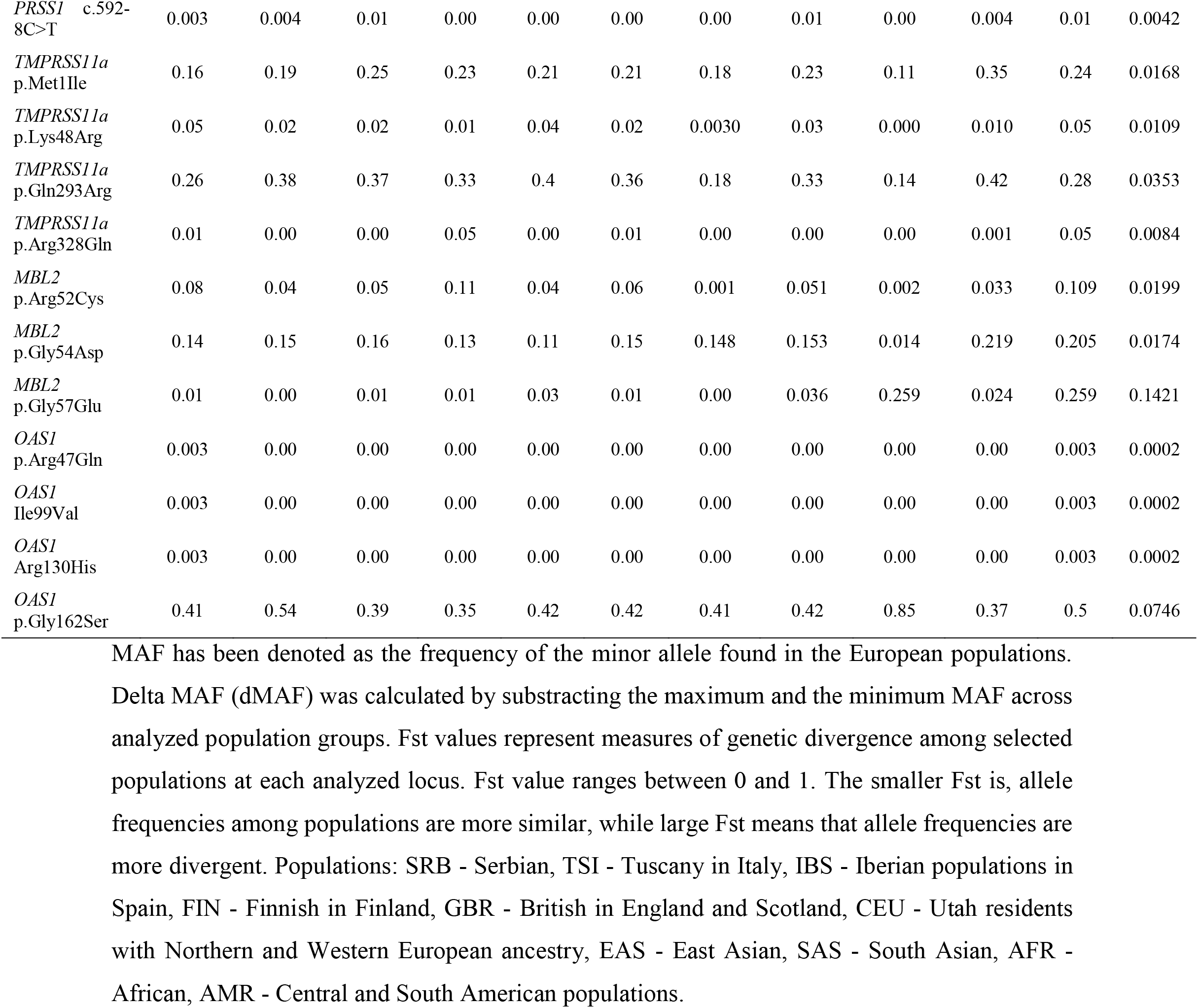
Minor allele frequencies (MAF) across Serbian and 1kGP populations.

The level of population genetic variability at each selected loci was assessed using two approaches, delta MAF and Fst statistics. Significant correlation between obtained delta MAF and Fst values (r=0.74, p=0.000033) was observed. MAF distributions of seven genetic variants (p.Asp472Asn in *PLG*; p.Met1Ile and p.Gln293Arg in *TMPRSS11a*; p.Arg52Cys, p.Gly54Asp and p.Gly57Glu in *MBL2*; p.Gly162Ser in *OAS1*), that demonstrated delta MAF>0.1 among analyzed populations, are presented in the Fig 1.

**Fig 1.**
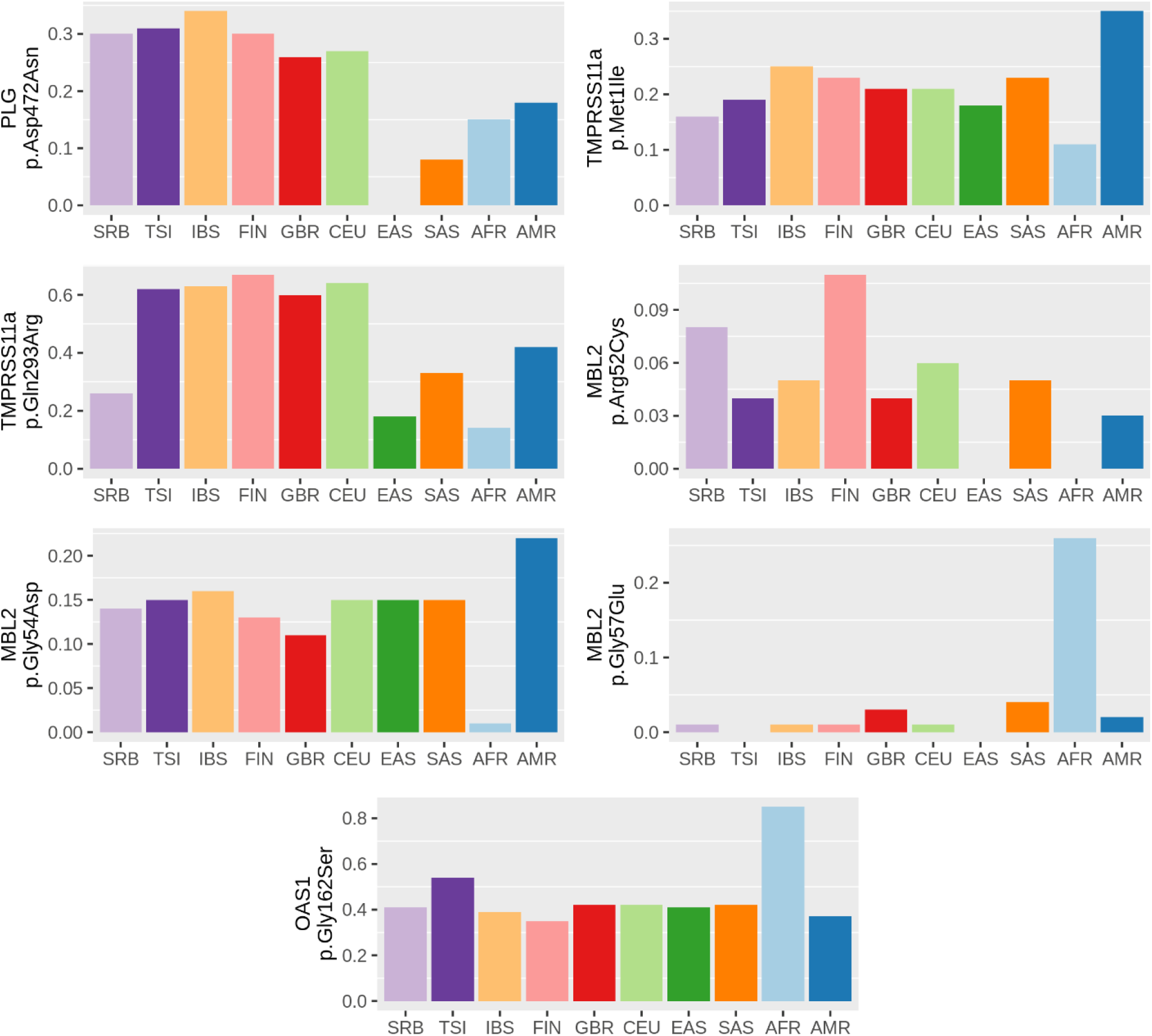
Distribution of MAF across Serbian and 1kGP populations. Distribution of MAF across Serbian and 1kGP populations for variants that showed highest genetic divergence (delta MAF>0.1). MAF value (0-1) of each genetic variant is presented on y axis. Populations: SRB - Serbian, TSI - Tuscany in Italy, IBS - Iberian populations in Spain, FIN - Finnish in Finland, GBR - British in England and Scotland, CEU - Utah residents with Northern and Western European ancestry, EAS - East Asian, SAS - South Asian, AFR - African, AMR - Central and South American populations.

Promisingly, three of those variants (p.Arg52Cys, p.Gly54Asp and p.Gly57Glu in *MBL2*), that showed considerable divergence in frequencies among all analyzed populations, were also predicted to have damaging effect on the protein. Variants p.Gly54Asp and p.Gly57Glu showed extreme MAF values in African compared to other populations (p.Gly54Asp MAF range: 1.4% in African to the 21.9% in Central and South American; p.Gly57Glu MAF range: 0% in Italian and East Asians to the 25.9% in African). *MBL2* p.Arg52Cys variant showed variable distribution among analyzed populations, having the highest MAF in Finish (11%), followed by Serbian population (8%), while the lowest MAF was in East Asian and African populations (0%).

Having in mind that variants in proteins involved in the innate immunity could be unfavorable in COVID-19 patients, further studies on variant *MBL2* p.Gly54Asp in Central and South American and on variant *MBL2* p.Gly57Glu in African populations are indicated.

Furthermore, for these seven most divergent variants, we calculated Fisher exact test statistics of Serbian against 1kGP populations based on their genotypic frequencies (Table 3). After Bonferoni correction, significant differences were observed in distribution of *PLG* p.Asp472Asn and *MBL2* p.Arg52Cys compared to Asian populations. In comparison with African populations, all variants, except *TMPRSS11a* p.Met1Ile, had significantly distinct distributions. Significant differences were observed as well compared to Central and South American populations in distribution of *PLG* p.Asp472Asn, *TMPRSS11a* p.Gln293Arg and p.Met1Ile variants.

**Table 3.**
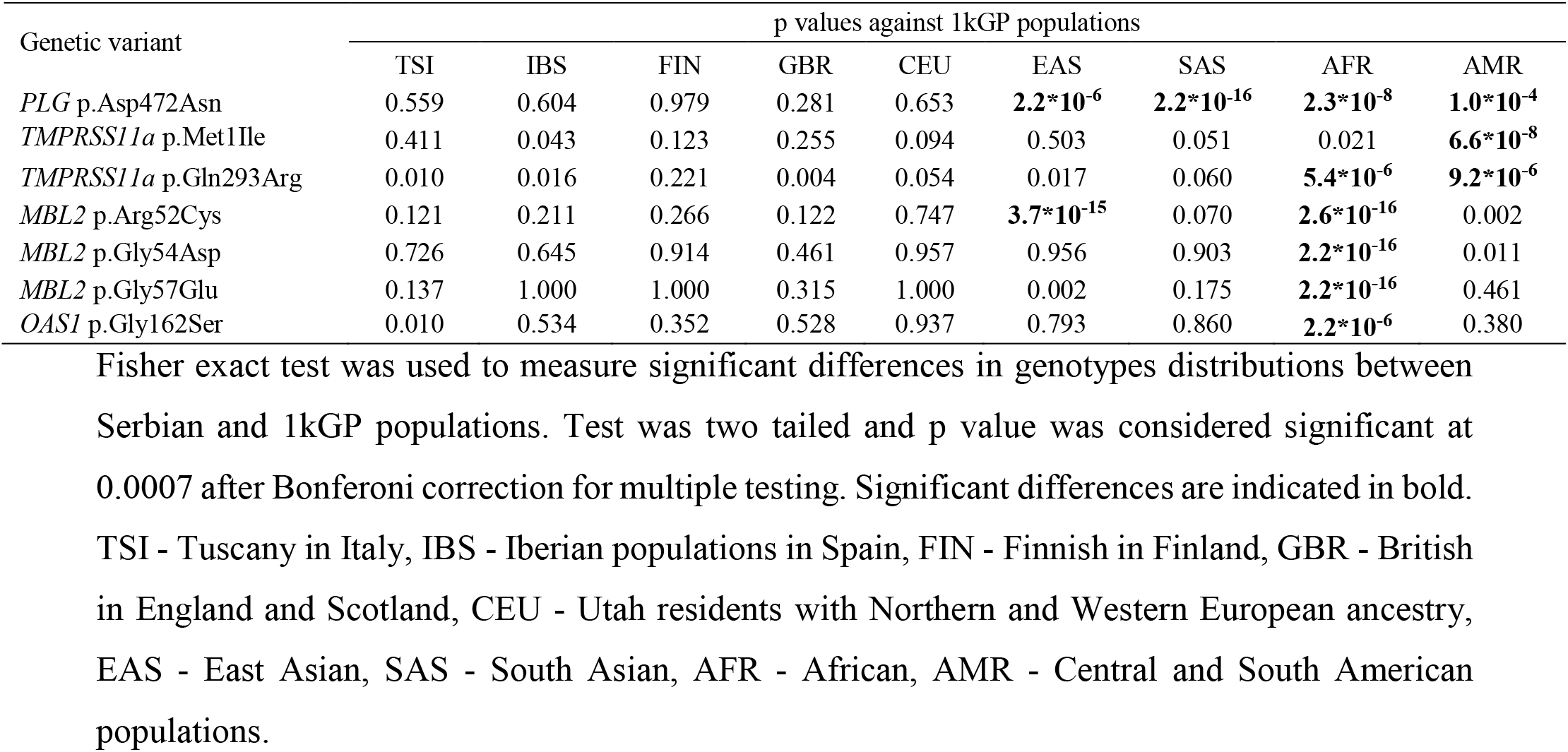
Comparison of Serbian with 1kGP populations in genotype distribution of selected variants.

| Genetic variant | p values against 1kGP populations |  |  |  |  |  |  |  |  |
| --- | --- | --- | --- | --- | --- | --- | --- | --- | --- |
|  | TSI | IBS | FIN | GBR | CEU | EAS | SAS | AFR | AMR |
| *PLG* p.Asp472Asn | 0.559 | 0.604 | 0.979 | 0.281 | 0.653 | **2.2\*10<sup>-6</sup>** | **2.2\*10<sup>-16</sup>** | **2.3\*10<sup>-8</sup>** | **1.0\*10<sup>-4</sup>** |
| *TMPRSS11a* p.Met1Ile | 0.411 | 0.043 | 0.123 | 0.255 | 0.094 | 0.503 | 0.051 | 0.021 | **6.6\*10<sup>-8</sup>** |
| *TMPRSS11a* p.Gln293Arg | 0.010 | 0.016 | 0.221 | 0.004 | 0.054 | 0.017 | 0.060 | **5.4\*10<sup>-6</sup>** | **9.2\*10<sup>-6</sup>** |
| *MBL2* p.Arg52Cys | 0.121 | 0.211 | 0.266 | 0.122 | 0.747 | **3.7\*10<sup>-15</sup>** | 0.070 | **2.6\*10<sup>-16</sup>** | 0.002 |
| *MBL2* p.Gly54Asp | 0.726 | 0.645 | 0.914 | 0.461 | 0.957 | 0.956 | 0.903 | **2.2\*10<sup>-16</sup>** | 0.011 |
| *MBL2* p.Gly57Glu | 0.137 | 1.000 | 1.000 | 0.315 | 1.000 | 0.002 | 0.175 | **2.2\*10<sup>-16</sup>** | 0.461 |
| *OAS1* p.Gly162Ser | 0.010 | 0.534 | 0.352 | 0.528 | 0.937 | 0.793 | 0.860 | **2.2\*10<sup>-6</sup>** | 0.380 |

Fisher exact test was used to measure significant differences in genotypes distributions between Serbian and 1kGP populations. Test was two tailed and p value was considered significant at 0.0007 after Bonferoni correction for multiple testing. Significant differences are indicated in bold. TSI - Tuscany in Italy, IBS - Iberian populations in Spain, FIN - Finnish in Finland, GBR - British in England and Scotland, CEU - Utah residents with Northern and Western European ancestry, EAS - East Asian, SAS - South Asian, AFR - African, AMR - Central and South American populations.

Finally, we analyzed genetic variation related to selected loci between Serbian and 1kGP populations using pairwise Fst calculation [25] (Table 4). Fst value ranges between 0 and 1. The smaller Fst indicates similar allele frequencies among populations, and vice versa, large Fst means that allele frequencies are more divergent. Comparing Serbian to other populations, the highest level of genetic differentiation related to selected loci was observed with African (Fst=0.147), followed by East Asian (Fst=0.054), Central and South American populations (0.036) and South Asian population (0.026). When compared with European populations, the highest divergence was observed with Italian population (Fst=0.012).

**Table 4.** Pairwise Fst values across Serbian and 1kGP populations.

|  | SRB | TSI | IBS | CEU | GBR | FIN | EAS | SAS | AFR | AMR |
| --- | --- | --- | --- | --- | --- | --- | --- | --- | --- | --- |
| SRB | NA |  |  |  |  |  |  |  |  |  |
| TSI | 0.012 | NA |  |  |  |  |  |  |  |  |
| IBS | 0.007 | 0.007 | NA |  |  |  |  |  |  |  |
| CEU | 0.003 | 0.003 | -0.001 | NA |  |  |  |  |  |  |
| GBR | 0.009 | 0.005 | 0.001 | -0.003 | NA |  |  |  |  |  |
| FIN | 0.004 | 0.016 | 0.000 | 0.000 | 0.004 | NA |  |  |  |  |
| EAS | 0.054 | 0.081 | 0.078 | 0.056 | 0.062 | 0.066 | NA |  |  |  |
| SAS | 0.026 | 0.033 | 0.030 | 0.016 | 0.016 | 0.025 | 0.020 | NA |  |  |
| AFR | 0.147 | 0.135 | 0.179 | 0.159 | 0.160 | 0.185 | 0.184 | 0.157 | NA |  |
| AMR | 0.036 | 0.031 | 0.015 | 0.015 | 0.015 | 0.019 | 0.064 | 0.018 | 0.202 | NA |

Fst values matrix showing genetic divergence between each population using genotype data for all analysed loci (22 selected variants). Fst value ranges between 0 and 1 (negative values are considered as zeros). The smaller Fst is, allele frequencies among populations are more similar, while large Fst means that allele frequencies are more divergent. Populations: SRB - Serbian, TSI - Tuscany in Italy, IBS - Iberian populations in Spain, FIN - Finnish in Finland, GBR - British in England and Scotland, CEU - Utah residents with Northern and Western European ancestry, EAS - East Asian, SAS - South Asian, AFR - African, AMR - Central and South America.

## Discussion

While mutations in the RNA of betacoronavirus opened the possibility to jump to a new specie, variants in various human genes may contribute to increased susceptibility to a new pathogen, or could have a protective role. One of the most illustrious examples are variants in *CD4* receptor which contribute to susceptibility to some strains of HIV infection, while variants in *CCR5* gene provide resistance to its carriers [27, 28].

Recent studies addressed rare variants in *ACE2* gene which could directly influence the binding to S protein of the SARS-CoV-2 and the population-specific differences of more frequent *ACE2* gene variants [21, 22]. However, having in mind complexity of the interaction between the virus and the host, which besides the S protein – ACE2 receptor interaction, includes the role of different host proteases and at least some elements of innate immunity with antiviral activity, in this study we selected additional genes that could be important for the susceptibility or resistance to the SARS-CoV-2 viral infection. We focused on genetic variants in genes *FURIN, PLG, PRSS1, TMPRSS11a,* that encode proteases, as well as variants in genes *MBL2* and *OAS1*, that encode proteins involved in innate immunity, attempting to find promising candidate alleles that might be included in the future genetic test related to new waives of the SARS-CoV-2 pandemics.

To systematically examine the coding variants in selected genes and differences in the allele frequency between populations, we performed comparative analysis of 22 variants found in *FURIN, PLG, PRSS1, TMPRSS11a, MBL2* and *OAS1* genes from Serbian population database with different European populations and super-populations extracted from 1kGP database [24]. Also, we used bioinformatic tools to predict the effect of these genetic variants on the structure and/or function of proteins encoded by *FURIN, PLG, PRSS1, TMPRSS11a, MBL2* and *OAS1* genes.

It could be possible that the existence of variants in genes encoding proteases provide advantage while variants in genes for proteins involved in the innate immunity add some disadvantage to individuals in combating COVID-19.

### Variants in genes for host proteases

Host proteases, such as furin, plasmin, trypsin-1 and TMPRSS11a, are crucial in the process of cutting the S protein of the SARS-CoV and SARS-CoV-2 envelope at the S1/S2 cleavage site, which is a necessary event needed to release the S1 fragment (N-terminal part) and the S2 fragment (C-terminal part) [17]. The S1 fragment recognizes ACE2 receptor at the surface of the human cells, and the S2 fragment is involved in the viral entry into the cells [29]. Furthermore, the spike protein of the SARS-CoV-2 must be cleaved at both, S1/S2 cleavage site and at the furin-like cleavage site inside the S2 fragment in order to enable viral entry [17]. It was previously shown that the cleavage by host proteases plasmin, trypsin-1 and TMPRSS11a at both sites of the S protein is mandatory for the entry of SARS-CoV into human bronchial epithelial cells *in vitro* [30]. Interestingly, the furin like S2’ cleavage site is identical between SARS-CoV and SARS-CoV-2 [17] thus implying the capability of furin, plasmin, trypsin-1 and TMPRSS11a to perform activation steps and enable viral entry into the human cells. The efficiency of all these four proteases contribute to regulation of cellular tropism and determination of viral pathogenesis. Thus, genetic variants that would impact the structure and/or function of furin, plasmin, trypsin-1 and TMPRSS11 may lead to inter-individual differences in the response to the SARS-CoV-2 infection.

*FURIN* encodes a type 1 membrane bound protease, one of the seven basic amino acid-specific members which cleave their substrates at single or paired basic residues. First, furin is autocatalyticaly processed in the endoplasmatic reticulum and then transported to the trans-Golgi network where a second autocatalytic event takes place and the catalytic activity is acquired. In addition to having affinity for various substrates in human tissues, it is probably one of the proteases responsible for the activation of SARS-CoV-2 envelope S glycoproteins [17]. Two rare variants, p.Thr33Ala and p.Gly146Ser, were detected in *FURIN*. Thr33 residue is located in the pro-domain of a peptidase and therefore cleaved off in the process of activation of the furin enzyme, whereas Gly146 residue is located within the peptidase domain of the enzyme [31]. A substitution of Gly146 to Ser is predicted to be probably damaging/deleterious by PolyPhen-2 and SIFT algorithms, while MutPred2 software has several hypotheses, including altered metal binding and altered catalytic site. Also, when we analyzed the effect of this variant upon the structure of furin protein in Swiss-Pdb Viewer, a change to Ser at position 146 is predicted to cause steric clashes with His145, which may lead to an unstable protein. Taken altogether, p.Gly146Ser variant may amend the activity of furin in its proprotein convertase function, and may change its ability to cleave furin-like sites in the S protein of the SARS-CoV-2.

Plasminogen is a protease which belongs to peptidase family S1 and it is encoded by the *PLG* gene. Proteolysis of plasminogen results in multiple forms of the active plasmin. Plasminogen is present in the human lung tissues, more precisely in the airway, alveolar type I and II epithelial cells as well as in endothelial cells [20]. Six very rare variants were identified in the *PLG* gene: p.Glu55Gln, p.Arg89Lys, p.Arg261His, p.Gly420Asp, detected only on one chromosome (MAF 0.003), and p.Ala494Val, p.Arg523Trp on two chromosomes (MAF 0.007). Variants p.Arg261His and p.Ala494Val are predicted to be probably damaging/deleterious, while p.Glu55Gln, p.Arg89Lys, p.Gly420Asp, and p.Arg523Trp are mostly predicted to be benign/tolerated by PolyPhen-2, SIFT and MutPred2 algorithms. Residues Glu55 and Arg89 are located in the Pan-Apple domain, while Arg261, Gly420A, Ala494 and Arg523 reside in different Kringle domains of the proenzyme precursor plasminogen [32]. Given that the Pan-Apple domains mediate protein-protein or protein-carbohydrate interactions, whereas Kringle domains play a role in binding mediators, these variants may be involved in fine tuning of plasmin activity. Having in mind that individuals with elevated plasmin had greater susceptibility to COVID-19 and more severe clinical manifestations [20], rare variants in *PLG*, such as p.Arg261His and p.Ala494Val may be recognized as potential markers of inter-individual differences in susceptibility to coronavirus.

Trypsin-1 is encoded by the *PRSS1* gene which is a member of the trypsin-1 family of serine proteases. It is active on peptide linkages involving the carboxyl group of lysine or arginine. This enzyme is secreted by the pancreas and cleaved to its active form in the small intestine. It is also present in airway and alveolar type I and II epithelial cells [20]. Two rare variants in the *PRSS1* gene, c.592-8C>T and p.Asn54Lys, were detected. Variant c.592-8C>T was previously detected in patients with cystic fibrosis presenting with chronic pancreatitis [33]. Given that this is a splice region variant, it may affect splicing and therefore the level of trypsin-1 protein. Trypsin-1 variant p.Asn54Lys leads to the substitution of polar asparagine residue to a positively charged lysine. This variant is predicted to be possibly damaging/deleterious by PolyPhen-2 and SIFT algorithms, while MutPred2 software suggested that this change may alter the catalytic site. Given that the catalytic triad of trypsin-1 enzyme is consisted of His57, Asn102 and Ser195 [34], the proximity of Asn54 to His57 might modify the enzymatic activity. The same codon is affected by an alternative variant, p.Asn54Ser, which was classified pathogenic by UniProt, and associated with chronic pancreatitis [35, 36]. Nevertheless, having in mind that the variant p.Asn54Lys was observed in healthy adults, this variant is not regarded as a disease causing in the context of Mendelian diseases. Its mechanism might be through the fine modification of trypsin-1 enzymatic activity, which could contribute to the resistance of some individuals to SARS-CoV-2 infection.

Transmembrane serine protease 11A, encoded by the TMPRSS11a gene is one of the members of type II transmembrane serine proteases and it is present in upper respiratory tract (pharynx and trachea) and digestive tract [30]. Two rare variants were detected in the *TMPRSS11A* gene: p.Lys48Arg and p.Arg328Gln. For both of these variants, PolyPhen-2, SIFT and MutPred2 algorithms predict benign/tolerated effect. Given the scarce data on the protein structure of *TMPRSS11A*, the effect of this variants is yet to be ascertained.

### Variants in genes for proteins involved in the host innate immunity

Innate immunity is important in defending organism from viral infections. After SARS-CoV outbreak it was found that type I interferons could inhibit the replication of this virus and that these interferons further induce different proteins with antiviral activity [37]. One of such proteins is encoded by the *OAS1* gene. OAS1 synthesizes 2’,5’-oligoadenylates, and as a consequence activates latent RNase L which cuts single-stranded RNAs thus leading to the viral RNA degradation and inhibition of viral replication [38]. Three rare variants were detected in the *OAS1* gene: p.Arg47Gln, p.Ile99Val and p.Arg130His. Interestingly, human OAS1 protein recognizes two adjacent minor grooves of dsRNA with C-terminal lobe and N-terminal lobe, where the Arg47 residue is one of the residues involved in the recognition [39]. When analyzed in Swiss-Pdb Viewer, the side chain of Arg47 residue was predicted to form hydrogen bond with dsRNA, while a change to glutamine at this position was predicted to abolish this hydrogen bond. Furthermore, Gln at this position is predicted to form hydrogen bonds with Cys45 and Phe46, and also to cause steric clashes with Glu43 and Arg44. Nevertheless, variant p.Arg47Gln is predicted to be benign/tolerated by PolyPhen-2, SIFT and MutPred2 algorithms, so its effect on the OAS1 protein seems not to affect the structure itself, but it may be apparent upon the binding of the dsRNA. Variant p.Ile99Val is predicted to be possibly damaging/deleterious by prediction algorithms, but the analysis in Swiss-Pdb Viewer revealed that the backbone of the Ile99 residue is predicted to form hydrogen bonds with Arg95, Gly96 and Arg103, so the change to Val at this position did not change the existing bonds, nor did it form novel ones. Variant p.Arg130His is predicted to be probably damaging/deleterious by prediction algorithms, and the analysis in Swiss-Pdb Viewer showed that in case of Arg130, a change to His at this position was predicted to result in the formation of novel hydrogen bonds with Thr188 and also a steric clash with Glu185. These rare variants may have an effect on the OAS1 structure and/or function through the formation of novel hydrogen bonds and steric clashes, leading to a less stable protein. In case of variants located at the protein/RNA interface formed upon the binding, its effects reflect on the formation of (to some extent) weaker bond with the RNA thus lowering its 2’-5’-oligoadenylate synthetase activity, as it was shown that variants at the protein/RNA interface impair OAS1 activity by 60- to 2,500-fold [39]. Furthermore, having in mind that the active form of human OAS1 is tetrameric, any variant potentially impairing the oligomerization process would also have an effect on the activity of the enzyme, the synthesis of 2’,5’-oligoadenylates, and consequently the activation of the latent RNase and the degradation of viral RNA of SARS-CoV-2. Knowing that variants in *OAS1*, such as p.Gly397Arg were previously associated with susceptibility to SARS-CoV infection in some populations [5], the effects of p.Arg47Gln, p.Ile99Val and p.Arg130His might be interesting for further investigation.

### Comparative population analysis

The comparative population analysis pointed to the 7 coding variants in *PLG, TMPRSS11a, MBL2* and *OAS1* with noticeable divergence in allelic frequencies between analyzed populations. Three variants were found in genes encoding proteases. Variant p.Asp472Asn in *PLG* gene showed notable MAF discrepancy between European and Asian populations, being the highest in Spain (0.34) and the lowest in East Asians (0.0). This substitution of neutral to acidic amino acid arises in a loop region that connects two Kringle domains of plasminogen and it was presumed to have a functional effect through altering the alignment of Kringle domains [40]. This variant was shown previously to influence susceptibility to invasive aspergillosis [40]. Having in mind the MAF difference between Spanish and East Asian populations, it would be interesting to further investigate whether it also reflects the remarkable differences in susceptibility and morbidity to COVID-19 between these populations. Two variants in *TMPRSS11A* showed interesting data for Serbian population, with both variants showing lower MAF in Serbian population (0.16 for p.Met1Ile and 0.26 for p.Gln293Arg) compared to other European populations - average MAF 0.22 for p.Met1Ile and 0.37 for p.Gln293Arg. As a start loss variant, p.Met1Ile leads to start of the translation from the neighboring Met2 residue. Start loss variants can range from disease causing to benign start codon variants, so with the high frequency of p.Met1Ile in all analyzed populations, its effect is presumed to be benign. While variant p.Gln293Arg was described as a cumulative risk factor for esophageal squamous cell carcinoma [41], its effect on the protease activity and COVID-19 susceptibility remains to be elucidated.

Four variants were found in genes involved in the innate immunity. Variant p.Gly162Ser in *OAS1* was shown to have highest MAF in African populations - 0.85, more than double the average MAF of European populations (0.42). This variant was previously weakly associated with type I diabetes [42] and multiple sclerosis [43], but a more recent study showed that this variant was found not to interfere with OAS1 enzyme activity [44].

Another important element of the innate immune system is mannose-binding protein (soluble mannose-binding lectin) which is encoded by the *MBL2* gene. The protein recognizes mannose and N-acetylglucosamine expressed on the surface of many microorganisms, and is capable of activating the classical complement pathway. Deficiencies of this gene have been associated with increased susceptibility to SARS-CoV and other autoimmune and infectious diseases [6]. Interestingly, out of 7 genetic variants that demonstrated delta MAF>0.1 in our study, only p.Arg52Cys, p.Gly54Asp and p.Gly57Glu in *MBL2* were predicted to be probably damaging/deleterious by prediction algorithms. Moreover, these three variants were already extensively studied and functionally characterized - they showed compromised oligomerization and thus the activity of the final protein [45]. In addition to promoter variants, these coding variants have been shown to influence the stability and serum concentration of the protein, where, consequently, low levels of MBL2 have been associated with increased susceptibility to infections [46]. Therefore, the influence of these three variants was studied in various immunological sceneries, from autoimmune to infections, including SARS-CoV [6, 47]. Nevertheless, all three variants are found in all analyzed populations, with relatively high MAF values. Interestingly, p.Gly54Asp and p.Gly57Glu showed extreme MAF values in African compared to other populations: p.Gly54Asp with MAF ranging from 0.014 in African to 0.219 in Central and South American and p.Gly57Glu with MAF ranging from 0.0 in Italian and East Asians to 0.259 in African. Variant p.Arg52Cys showed variable distribution among analyzed populations, having the highest MAF in Finish (0.11), followed by Serbian population (0.08), while the lowest MAF was in East Asian and African populations (0.0). Further studies in the populations in which those variants are frequent could contribute to design of prediction model of the SARS-CoV-2 susceptibility in the carriers of the variants.

Taken altogether, comparing Serbian to other populations, it was found that the highest level of genetic differentiation related to selected loci was observed with African, followed by East Asian and South Asian populations. When compared with European populations, the highest divergence was observed with Italian population.

In conclusion, results of our analysis showed that variants predicted to have altering effect to the proteins are very rare in each of the selected European populations as well as super-populations. Thus, although it is not likely to perform massive genetic testing aiming to detect these variants in order to predict prognosis to COVID-19, they may provide answers for the inter-individual differences in the clinical course of disease among patients of the same age and the same genetic background which received identical medical treatment.

On the other hand, variants which have divergent allele frequencies between populations were mostly predicted to lack effect on the structure and/or the function of the proteins. However, few variants, such as those in *MBL2* gene were predicted to have some functional effect and their contribution to population differences regarding COVID-19 should be further evaluated.

In general, findings of this study may lead us to conclude that 4 coding variants in genes encoding proteases (*FURIN, PLG* and *PRSS1*) and 6 in genes involved in the innate immunity (*MBL2* and *OAS1*) could be considered as candidates in forthcoming studies aiming to explain differences in clinical manifestations, recovery rate and mortality rate of COVID-19 which vary between different populations [48]. Further genetic analysis of variants in the non-coding regions (contributing to the fine tuning of the gene expression) of these genes, as well as variants in other genes, are needed to complement our study and give the complete insight about inter-individual and population-specific genetic susceptibility and resistance to the SARS-CoV-2 infection. A global genetic initiative [49] holds promise that the full spectrum of human genetic factors determining susceptibility, severity and outcomes of COVID-19 will be determined in future.

## Acknowledgments

This work was supported by Ministry of Education, Science and Technological Development Republic of Serbia, EB: 451-03-68/2020-14/ 200042.

## References

1. Wu F, Zhao S, Yu B, Chen YM, Wang W, Song ZG, et al. A new coronavirus associated with human respiratory disease in China. Nature. 2020;579(7798):265–9. doi: 10.1038/s41586-020-2008-3. PubMed PMID: 32015508; PubMed Central PMCID: PMC7094943.

2. WHO. Available from: https://www.who.int/docs/default-source/coronaviruse/situation-reports/20200311-sitrep-51-covid-19.pdf?sfvrsn=1ba62e57_10.

3. Kenney AD, Dowdle JA, Bozzacco L, McMichael TM, St Gelais C, Panfil AR, et al. Human Genetic Determinants of Viral Diseases. Annual review of genetics. 2017;51:241–63. doi: 10.1146/annurev-genet-120116-023425. PubMed PMID: 28853921; PubMed Central PMCID: PMC6038703.

4. Williams FM, Freydin M, Mangino M, Couvreur S, Visconti A, Bowyer RC, et al. Self-reported symptoms of covid-19 including symptoms most predictive of SARS-CoV-2 infection, are heritable. medRxiv. 2020:2020.04.22.20072124. doi: 10.1101/2020.04.22.20072124.

5. He J, Feng D, de Vlas SJ, Wang H, Fontanet A, Zhang P, et al. Association of SARS susceptibility with single nucleic acid polymorphisms of OAS1 and MxA genes: a case-control study. BMC infectious diseases. 2006;6:106. doi: 10.1186/1471-2334-6-106. PubMed PMID: 16824203; PubMed Central PMCID: PMC1550407.

6. Tu X, Chong WP, Zhai Y, Zhang H, Zhang F, Wang S, et al. Functional polymorphisms of the CCL2 and MBL genes cumulatively increase susceptibility to severe acute respiratory syndrome coronavirus infection. The Journal of infection. 2015;71(1):101–9. doi: 10.1016/j.jinf.2015.03.006. PubMed PMID: 25818534; PubMed Central PMCID: PMC7112636.

7. Zhu X, Wang Y, Zhang H, Liu X, Chen T, Yang R, et al. Genetic variation of the human alpha-2-Heremans-Schmid glycoprotein (AHSG) gene associated with the risk of SARS-CoV infection. PloS one. 2011;6(8):e23730. doi: 10.1371/journal.pone.0023730. PubMed PMID: 21904596; PubMed Central PMCID: PMC3163911.

8. Ng MW, Zhou G, Chong WP, Lee LW, Law HK, Zhang H, et al. The association of RANTES polymorphism with severe acute respiratory syndrome in Hong Kong and Beijing Chinese. BMC infectious diseases. 2007;7:50. doi: 10.1186/1471-2334-7-50. PubMed PMID: 17540042; PubMed Central PMCID: PMC1899505.

9. Chong WP, Ip WK, Tso GH, Ng MW, Wong WH, Law HK, et al. The interferon gamma gene polymorphism +874 A/T is associated with severe acute respiratory syndrome. BMC infectious diseases. 2006;6:82. doi: 10.1186/1471-2334-6-82. PubMed PMID: 16672072; PubMed Central PMCID: PMC1468415.

10. Yuan FF, Boehm I, Chan PK, Marks K, Tang JW, Hui DS, et al. High prevalence of the CD14-159CC genotype in patients infected with severe acute respiratory syndrome-associated coronavirus. Clinical and vaccine immunology : CVI. 2007;14(12):1644–5. doi: 10.1128/CVI.00100-07. PubMed PMID: 17913858; PubMed Central PMCID: PMC2168372.

11. Chan KY, Xu MS, Ching JC, Chan VS, Ip YC, Yam L, et al. Association of a single nucleotide polymorphism in the CD209 (DC-SIGN) promoter with SARS severity. Hong Kong medical journal = Xianggang yi xue za zhi. 2010;16(5 Suppl 4):37–42. PubMed PMID: 20864747.

12. Lu G, Wang Q, Gao GF. Bat-to-human: spike features determining ‘host jump’ of coronaviruses SARS-CoV, MERS-CoV, and beyond. Trends in microbiology. 2015;23(8):468–78. doi: 10.1016/j.tim.2015.06.003. PubMed PMID: 26206723; PubMed Central PMCID: PMC7125587.

13. Chan JF, Kok KH, Zhu Z, Chu H, To KK, Yuan S, et al. Genomic characterization of the 2019 novel human-pathogenic coronavirus isolated from a patient with atypical pneumonia after visiting Wuhan. Emerging microbes & infections. 2020;9(1):221–36. doi: 10.1080/22221751.2020.1719902. PubMed PMID: 31987001; PubMed Central PMCID: PMC7067204.

14. Wrapp D, Wang N, Corbett KS, Goldsmith JA, Hsieh CL, Abiona O, et al. Cryo-EM structure of the 2019-nCoV spike in the prefusion conformation. Science. 2020;367(6483):1260–3. doi: 10.1126/science.abb2507. PubMed PMID: 32075877; PubMed Central PMCID: PMC7164637.

15. Matsuyama S, Nagata N, Shirato K, Kawase M, Takeda M, Taguchi F. Efficient activation of the severe acute respiratory syndrome coronavirus spike protein by the transmembrane protease TMPRSS2. Journal of virology. 2010;84(24):12658–64. doi: 10.1128/JVI.01542-10. PubMed PMID: 20926566; PubMed Central PMCID: PMC3004351.

16. Hoffmann M, Kleine-Weber H, Schroeder S, Kruger N, Herrler T, Erichsen S, et al. SARS-CoV-2 Cell Entry Depends on ACE2 and TMPRSS2 and Is Blocked by a Clinically Proven Protease Inhibitor. Cell. 2020;181(2):271–80 e8. doi: 10.1016/j.cell.2020.02.052. PubMed PMID: 32142651; PubMed Central PMCID: PMC7102627.

17. Coutard B, Valle C, de Lamballerie X, Canard B, Seidah NG, Decroly E. The spike glycoprotein of the new coronavirus 2019-nCoV contains a furin-like cleavage site absent in CoV of the same clade. Antiviral research. 2020;176:104742. doi: 10.1016/j.antiviral.2020.104742. PubMed PMID: 32057769; PubMed Central PMCID: PMC7114094.

18. Chen J, Lee KH, Steinhauer DA, Stevens DJ, Skehel JJ, Wiley DC. Structure of the hemagglutinin precursor cleavage site, a determinant of influenza pathogenicity and the origin of the labile conformation. Cell. 1998;95(3):409–17. doi: 10.1016/s0092-8674(00)81771-7. PubMed PMID: 9814710.

19. Zhao R, Ali G, Nie HG, Chang Y, Bhattarai D, Su X, et al. Plasmin improves oedematous blood-gas barrier by cleaving epithelial sodium channels. British journal of pharmacology. 2020. doi: 10.1111/bph.15038. PubMed PMID: 32133621.

20. Ji HL, Zhao R, Matalon S, Matthay MA. Elevated Plasmin(ogen) as a Common Risk Factor for COVID-19 Susceptibility. Physiological reviews. 2020;100(3):1065–75. doi: 10.1152/physrev.00013.2020. PubMed PMID: 32216698.

21. Hussain M, Jabeen N, Raza F, Shabbir S, Baig AA, Amanullah A, et al. Structural variations in human ACE2 may influence its binding with SARS-CoV-2 spike protein. Journal of medical virology. 2020. doi: 10.1002/jmv.25832. PubMed PMID: 32249956.

22. Cao Y, Li L, Feng Z, Wan S, Huang P, Sun X, et al. Comparative genetic analysis of the novel coronavirus (2019-nCoV/SARS-CoV-2) receptor ACE2 in different populations. Cell discovery. 2020;6:11. doi: 10.1038/s41421-020-0147-1. PubMed PMID: 32133153; PubMed Central PMCID: PMC7040011.

23. Skakic A, Djordjevic M, Sarajlija A, Klaassen K, Tosic N, Kecman B, et al. Genetic characterization of GSD I in Serbian population revealed unexpectedly high incidence of GSD Ib and 3 novel SLC37A4 variants. Clinical genetics. 2018;93(2):350–5. doi: 10.1111/cge.13093. PubMed PMID: 28685844.

24. Genomes Project C, Auton A, Brooks LD, Durbin RM, Garrison EP, Kang HM, et al. A global reference for human genetic variation. Nature. 2015;526(7571):68–74. doi: 10.1038/nature15393. PubMed PMID: 26432245; PubMed Central PMCID: PMC4750478.

25. Nei M. Molecular Evolutionary Genetics: Columbia University Press; 1987.

26. Holsinger KE, Weir BS. Genetics in geographically structured populations: defining, estimating and interpreting F(ST). Nature reviews Genetics. 2009;10(9):639–50. doi: 10.1038/nrg2611. PubMed PMID: 19687804; PubMed Central PMCID: PMC4687486.

27. Marmor M, Hertzmark K, Thomas SM, Halkitis PN, Vogler M. Resistance to HIV infection. Journal of urban health : bulletin of the New York Academy of Medicine. 2006;83(1):5–17. doi: 10.1007/s11524-005-9003-8. PubMed PMID: 16736351; PubMed Central PMCID: PMC1539443.

28. Oyugi JO, Vouriot FC, Alimonti J, Wayne S, Luo M, Land AM, et al. A common CD4 gene variant is associated with an increased risk of HIV-1 infection in Kenyan female commercial sex workers. The Journal of infectious diseases. 2009;199(9):1327–34. doi: 10.1086/597616. PubMed PMID: 19301975.

29. Wan Y, Shang J, Graham R, Baric RS, Li F. Receptor Recognition by the Novel Coronavirus from Wuhan: an Analysis Based on Decade-Long Structural Studies of SARS Coronavirus. Journal of virology. 2020;94(7). doi: 10.1128/JVI.00127-20. PubMed PMID: 31996437; PubMed Central PMCID: PMC7081895.

30. Kam YW, Okumura Y, Kido H, Ng LF, Bruzzone R, Altmeyer R. Cleavage of the SARS coronavirus spike glycoprotein by airway proteases enhances virus entry into human bronchial epithelial cells in vitro. PloS one. 2009;4(11):e7870. doi: 10.1371/journal.pone.0007870. PubMed PMID: 19924243; PubMed Central PMCID: PMC2773421.

31. Dahms SO, Arciniega M, Steinmetzer T, Huber R, Than ME. Structure of the unliganded form of the proprotein convertase furin suggests activation by a substrate-induced mechanism. Proceedings of the National Academy of Sciences of the United States of America. 2016;113(40):11196–201. doi: 10.1073/pnas.1613630113. PubMed PMID: 27647913; PubMed Central PMCID: PMC5056075.

32. Law RH, Caradoc-Davies T, Cowieson N, Horvath AJ, Quek AJ, Encarnacao JA, et al. The X-ray crystal structure of full-length human plasminogen. Cell reports. 2012;1(3):185–90. doi: 10.1016/j.celrep.2012.02.012. PubMed PMID: 22832192.

33. Sofia VM, Surace C, Terlizzi V, Da Sacco L, Alghisi F, Angiolillo A, et al. Trans-heterozygosity for mutations enhances the risk of recurrent/chronic pancreatitis in patients with Cystic Fibrosis. Molecular medicine. 2018;24(1):38. doi: 10.1186/s10020-018-0041-6. PubMed PMID: 30134826; PubMed Central PMCID: PMC6062922.

34. Polgar L. The catalytic triad of serine peptidases. Cellular and molecular life sciences : CMLS. 2005;62(19-20):2161–72. doi: 10.1007/s00018-005-5160-x. PubMed PMID: 16003488.

35. Teich N, Nemoda Z, Kohler H, Heinritz W, Mossner J, Keim V, et al. Gene conversion between functional trypsinogen genes PRSS1 and PRSS2 associated with chronic pancreatitis in a six-year-old girl. Human mutation. 2005;25(4):343–7. doi: 10.1002/humu.20148. PubMed PMID: 15776435; PubMed Central PMCID: PMC2752332.

36. Richards S, Aziz N, Bale S, Bick D, Das S, Gastier-Foster J, et al. Standards and guidelines for the interpretation of sequence variants: a joint consensus recommendation of the American College of Medical Genetics and Genomics and the Association for Molecular Pathology. Genetics in medicine : official journal of the American College of Medical Genetics. 2015;17(5):405–24. doi: 10.1038/gim.2015.30. PubMed PMID: 25741868; PubMed Central PMCID: PMC4544753.

37. Cinatl J, Morgenstern B, Bauer G, Chandra P, Rabenau H, Doerr HW. Treatment of SARS with human interferons. Lancet. 2003;362(9380):293–4. doi: 10.1016/s0140-6736(03)13973-6. PubMed PMID: 12892961; PubMed Central PMCID: PMC7112413.

38. Rebouillat D, Hovanessian AG. The human 2’,5’-oligoadenylate synthetase family: interferon-induced proteins with unique enzymatic properties. Journal of interferon & cytokine research : the official journal of the International Society for Interferon and Cytokine Research. 1999;19(4):295–308. doi: 10.1089/107999099313992. PubMed PMID: 10334380.

39. Donovan J, Dufner M, Korennykh A. Structural basis for cytosolic double-stranded RNA surveillance by human oligoadenylate synthetase 1. Proceedings of the National Academy of Sciences of the United States of America. 2013;110(5):1652–7. doi: 10.1073/pnas.1218528110. PubMed PMID: 23319625; PubMed Central PMCID: PMC3562804.

40. Zaas AK, Liao G, Chien JW, Weinberg C, Shore D, Giles SS, et al. Plasminogen alleles influence susceptibility to invasive aspergillosis. PLoS genetics. 2008;4(6):e1000101. doi: 10.1371/journal.pgen.1000101. PubMed PMID: 18566672; PubMed Central PMCID: PMC2423485 is a consultant for the Robert Michael Educational Institute. Guochun Liao, Jonathan Usuka and Gary Peltz are employees of Roche Biosciences. All other authors declare that no conflict of interest exists.

41. Suo C, Qing T, Liu Z, Yang X, Yuan Z, Yang YJ, et al. Differential Cumulative Risk of Genetic Polymorphisms in Familial and Nonfamilial Esophageal Squamous Cell Carcinoma. Cancer epidemiology, biomarkers & prevention : a publication of the American Association for Cancer Research, cosponsored by the American Society of Preventive Oncology. 2019;28(12):2014–21. doi: 10.1158/1055-9965.EPI-19-0484. PubMed PMID: 31562207.

42. Qu HQ, Polychronakos C, Type IDGC. Reassessment of the type I diabetes association of the OAS1 locus. Genes and immunity. 2009;10 Suppl 1:S69–73. doi: 10.1038/gene.2009.95. PubMed PMID: 19956105; PubMed Central PMCID: PMC2805449.

43. Fedetz M, Matesanz F, Caro-Maldonado A, Fernandez O, Tamayo JA, Guerrero M, et al. OAS1 gene haplotype confers susceptibility to multiple sclerosis. Tissue antigens. 2006;68(5):446–9. doi: 10.1111/j.1399-0039.2006.00694.x. PubMed PMID: 17092260.

44. Kjaer KH, Pahus J, Hansen MF, Poulsen JB, Christensen EI, Justesen J, et al. Mitochondrial localization of the OAS1 p46 isoform associated with a common single nucleotide polymorphism. BMC cell biology. 2014;15:33. doi: 10.1186/1471-2121-15-33. PubMed PMID: 25205466; PubMed Central PMCID: PMC4165621.

45. Larsen F, Madsen HO, Sim RB, Koch C, Garred P. Disease-associated mutations in human mannose-binding lectin compromise oligomerization and activity of the final protein. The Journal of biological chemistry. 2004;279(20):21302–11. doi: 10.1074/jbc.M400520200. PubMed PMID: 14764589.

46. Garred P, Larsen F, Seyfarth J, Fujita R, Madsen HO. Mannose-binding lectin and its genetic variants. Genes and immunity. 2006;7(2):85–94. doi: 10.1038/sj.gene.6364283. PubMed PMID: 16395391.

47. Ip WK, Chan KH, Law HK, Tso GH, Kong EK, Wong WH, et al. Mannose-binding lectin in severe acute respiratory syndrome coronavirus infection. The Journal of infectious diseases. 2005;191(10):1697–704. doi: 10.1086/429631. PubMed PMID: 15838797.

48. worldometers.info/coronavirus. Available from: worldometers.info/coronavirus.

49. www.covid19hg.org. Available from: www.covid19hg.org.

